# Effects of biological and environmental factors on orthoebolavirus serology in bats in Guinea

**DOI:** 10.1101/2023.08.23.554395

**Authors:** Maëliss Champagne, Julien Cappelle, Alexandre Caron, Thibault Pouliquen, Aboubacar Samoura, Mohamed Idriss Doumbouya, Guillaume Thaurignac, Abdoul Karim Soumah, Ahidjo Ayouba, Alpha Kabinet Keita, Martine Peeters, Mathieu Bourgarel, Hélène M. De Nys

**Author notes:** Correspondence should be addressed to Hélène M. De Nys.

## Abstract

**Background:** Outbreaks of Ebola disease emerge regularly in human populations on the African continent. This highly virulent haemorrhagic fever can be caused by different orthoebolaviruses from the *Filoviridae* family. Several bat species are believed to carry and transmit some of these viruses to people or other animal species. However, clear evidence is still lacking. Natural circulation and zoonotic transmissions of viruses are generally complex as they often involve several host species and are influenced by biological, environmental and anthropogenic factors. A better understanding of the role of bats and other wildlife in the emergence of Ebola disease outbreaks is crucial for their prevention and early detection.

**Methods:** In this longitudinal study we searched for antibodies against orthoebolaviruses in fruit bats in order to explore some of the factors which might influence their natural circulation. We performed serological tests on populations of 4 bat species (*Eidolon helvum*, *Hypsignathus monstrosus*, *Myonycteris angolensis* and *Rousettus aegyptiacus*) sampled longitudinally for 18 months (2018-2020) in Guinea. The analysis of 1,427 bat samples for antibodies directed against different orthoebolavirus species allowed us to test the influence of biological and environmental variables on seropositivity by using GLMM.

**Results:** Results showed that the presence of antibodies against one of the orthoebolaviruses antigen is more frequent in the *Eidolon helvum* and *Rousettus aegyptiacus* species, in males, in sexually immature adults, and during the dry season.

**Conclusion:** Our results suggest that host species, sex, age, reproductive life-cycle and season may play an important role in the circulation of filoviruses. This information can be used to guide further sampling to specifically characterize the orthoebolavirus species underlying the production of these antibodies and to adjust surveillance protocols at the interface between bats, humans and other animals.

## Background

Since the first two outbreaks of Ebola disease (EBOD) in 1976 in Yambuku (Democratic Republic of Congo) and Nzara (South Sudan), more than 30 outbreaks have been officially identified in humans across Central and West Africa (1). The frequency of these outbreaks seems to have recently increased (1).

Despite recent epidemics which originated from viral resurgence in human populations (2–4), most outbreaks are believed to have resulted from independent zoonotic transmissions, through direct or indirect contact with a putative animal reservoir or other infected animals (5). The reservoir species of viruses from the *Orthoebolavirus* genus remain unknown, but bats are suspected to play an important role in the ecology of these viruses and transmission mechanisms (6–8). It is hypothesized that bats might infect humans directly when hunted and handled (9), or indirectly through contact with bat excrement or fruit contaminated with saliva or excrement (10). Bats might also be an indirect source by transmitting viruses to bridge hosts which can act as amplifiers and be at the origin of human outbreaks (6,11). This has been suspected in Central Africa and Ivory Coast where apes have been confirmed as the source of human infection in several outbreaks (12,13).

Certain bat species are regarded as potential reservoir of orthoebolaviruses since the discovery of viral RNA and anti-Ebola virus (anti-EBOV) antibodies in 3 fruit bat species (*Epomops franqueti, Hypsignathus monstrosus, Myonycteris torquata*) (6). However, viral RNA was only detected in 13 of 279 tested specimens, and confirmed by sequence analysis in 7. Other studies have since shown that other frugivorous species have anti-orthoebolavirus antibodies (*Rousettus aegyptiacus, Eidolon helvum, Epomophorus gambianus, Micropteropus pusillus and Myonycteris angolensis*) and that insectivorous species (*Mops condylurus and Chaerephon pumilus*) host a recently discovered *Orthoebolavirus* named Bombali virus (7,8,14). This supports the fact that these wild bats are exposed to filoviruses and are able to survive infection, as demonstrated by experimental inoculations for some of these species (15,16). In addition, *Rousettus aegyptiacus* is the reservoir species of Marburg virus, another filovirus (17), and other bat species have been shown to be exposed or infected with other filoviruses such as Lloviu virus detected in Europe (18) and newly described viruses in China (19). Infectious orthoebolaviruses have never been isolated from free ranging bats so far. The viremic period might be short or viruses might end up circulating at a low level and might only be detected for a limited period of time in tissues or oral and rectal swabs, as shown experimentally in *R. aegyptiacus* inoculated with Ebola and Marburg virus (15,20).

The aerial lifestyle, gregariousness, longevity as well as immunological and metabolic specificities of bats are traits that may impact the ecology of orthoebolaviruses. Biological and phenological factors, influenced by seasonal or climate changes, are also important, like the migratory nature of certain bat species or reproduction patterns, both influenced by the availability of food resources (21,22). It has been shown that spillover events of Marburg virus to human populations coincide with the birthing season and presence of juveniles in *Rousettus aegyptiacus* populations (23). A recent study on an *Eidolon helvum* colony also showed an increase in the presence of antibodies against orthoebolaviruses when juveniles turn into immature adults (24). Other factors can influence the dynamics of orthoebolaviruses in their natural hosts, for instance population size and geographic distribution seem to impact viral richness within bat populations (25). All these factors influence to some extent viral circulation, risk of spillover events and emergence and are important to understand to adjust orthoebolavirus research and EBOD surveillance strategies.

The objective of this study was to investigate the effects of biological and environmental factors on the presence of anti-orthoebolavirus antibodies and improve understanding of dynamics of orthoebolavirus circulation in bats. We performed a longitudinal serological survey of orthoebolavirus in four frugivorous bat species in Guinea, a country at risk of EBOD outbreak. Based on the outcomes of the serological investigation, a subset of bats was also screened for filovirus RNA.

## Methods

### Sample and data collection

We collected blood samples as well as oral and rectal swabs from four frugivorous bat species, i.e. *Eidolon helvum (E. helvum), Hypsignathus monstrosus (H. monstrosus), Myonycteris angolensis (M. angolensis) and Rousettus aegyptiacus (R. aegyptiacus)*, in which anti-orthoebolavirus antibodies have already been detected in the past (8). Capture and sampling were carried out longitudinally over 13 sampling periods (i.e. 16 to 28 consecutive days of sampling per period) between November 2018 and July 2020, following the protocols already described in De Nys *et al.* (8). Bats were released directly after sampling. They were captured in feeding and roosting areas located mainly in four study sites of the prefecture of Macenta (Figure 1), in forested Guinea, the region where the 2013-2016 Ebola epidemic emerged. Sites were visited every 2 to 4 weeks during the course of the study. The environment was mainly constituted of highly transformed habitat, with secondary and degraded forests being gradually converted into a mosaic of forests and crops over the last 20 years (L. Lee-Cruz, pers. Comm.). It varied from secondary forest, fields, villages to more urbanised areas. Climate is characterized by a wet (May to October) and a dry (November to April) season.

**Figure 1.**
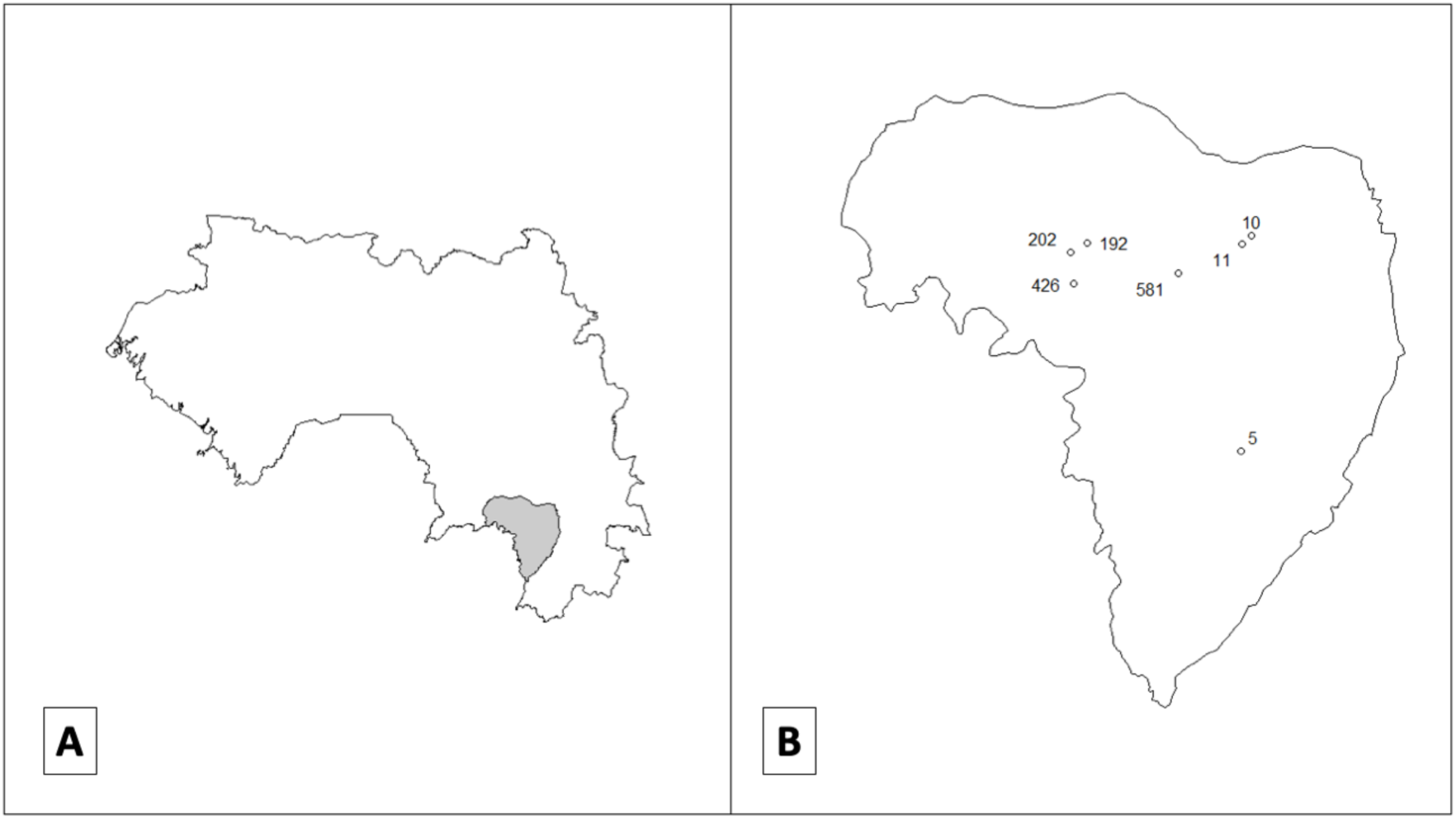
Sampling sites, A: Map of Guinea with the prefecture of Macenta in grey; B: prefecture of Macenta with sampling sites (dots) and number of sampled bats. Source of the basemap: https://data.humdata.org/dataset/cod-ab-gin; license: https://data.humdata.org/faqs/licenses.

Capture site (geographic coordinates, ecological environment), morphology, sex, age class, status of lactation and late gestation, and species (determined according to morphological criteria (26,27) and combined to genetic confirmation for some specimens, Additional file 1 Methods S1) were also recorded. Age was categorised as follows: juvenile (articular epiphyses not yet fully ossified), sexually immature adult (articular epiphyses ossified, but underdeveloped sexual organs), sexually mature adult (articular epiphyses ossified and clearly developed sexual organs).

This study was approved by the Comité National d’Ethique pour la Recherche en Santé of Guinea (permits 011/CNERS/19; 025/CNERS/20; 019/CNERS/20) and the Office Guinéen des Parcs et Réserves (permit 017/OGUIPAR/MEEF/2017).

### Screening for Ebolaviruses Antibodies

We tested dried blood spots (DBS) for anti-orthoebolavirus IgG with a Luminex-based serologic assay adapted for bats as described in De Nys *et al*. (8). The assay included ten recombinant orthoebolavirus proteins, i.e. the glycoprotein (GP), nucleoprotein (NP), and/or viral protein 40 (VP40), for different orthoebolaviruses: Ebola virus Kissidougou and Mayinga strains (NP, GP K, GP M, VP40), Sudan virus (NP, GP, VP40), Bundibugyo virus (GP, VP40) and Reston virus (GP). Recombinant proteins were produced in sf9 insect cell vectors and purchased at Sinobiologicals (Beijing, China) except for REST GP (IBT (Gaithersburg, MD)). The characteristics of the antigens of the different orthoebolavirus species used are as follows: Ebola virus (EBOV) NP (amino acids [aa] 488 to 739 strain Mayinga 1976), VP40 (aa 31 to 326, strain Kissidougou-Makona 2014), GP K (aa 1 to 650, Kissidougou-Makona 2014), GP M (aa 1 to 650, Mayinga 1976); Sudan virus (SUDV) NP (aa 361-738, strain Gulu), GP (aa 1 to 637, strain Uganda 2000), and VP40 (aa 31 to 326, strain Gulu); Bundibugyo virus (BDBV) GP (aa 1 to 501, strain Uganda 2007), and VP40 (aa 31 to 326, strain Uganda 2007); and Reston virus (RESTV) GP (aa 1 to 650). DBS were eluted and tested as previously described in 200µL of incubation buffer (8,24,28,29). One hundred µL of sample, corresponding to the equivalent of a final plasma dilution of 1/2000, was incubated with 50µL of magnetic beads coated with recombinant protein (2 μg protein/1.25 × 106 beads) in 96-well flat-bottom chimney plates (Greiner bio one, Frickenhausen, Germany) on a plate shaker at 300 rpm for 16 h at 4 °C in the dark. After washing, 0.1 μg/mL of goat anti-bat biotin-labeled IgG (Euromedex, Souffelweyersheim, France) was added to each well and incubated for 30 min at 300 rpm at room temperature. After washing, we added 50 μL of 4 μg/mL streptavidin-R-phycoerythrin (Fisher Scientific/Life Technologies, Illkirch, France) per well and incubated for 10 min at 300 rpm at room temperature. Reactions were read with BioPlex-200 (BioRad, Marnes-la-Coquette, France) or MagPix (Luminex, Austin, TX, US). At least 100 events were read for each bead set on a Bioplex-200 instrument, and results were expressed as median fluorescence intensity (MFI) per 100 beads. Samples that showed positive signals were repeated in order to validate the results.

To interpret the serological results in the absence of true positive controls, MFI cutoff values for each antigen were determined using four methods in De Nys et al. (8) (See Table S1 in Additional file 1). In this study we opted for the cutoff values resulting from the least stringent method, i.e. the mean plus four times the standard deviation of MFIs on negative control samples (M + 4SD) as calculated in De Nys et al. 2018, to search for serological patterns through a multivariate statistical analysis. Negative control samples (n = 145) were from captive-born insectivorous bat species (*Carollia perspicillata*, n=103) hosted at the Parc Zoologique de Montpellier (Montpellier, France) and two frugivorous bat species (*Pteropus giganteus*, n=19, *Rousettus aegyptiacus*, n=23) hosted at Wilhelma Zoo and Botanical Garden (Stuttgart, Germany). Samples were considered seropositive for an antigen if the MFI values were above the cutoff value for this antigen. Samples were considered seropositive for an orthoebolavirus if MFI values were above the cutoff value for at least two antigens of this specific orthoebolavirus.

### Multivariate statistical analyses

Proportions of samples positive to at least one orthoebolavirus antigen or to minimum two antigens of the same orthoebolavirus are presented in a descriptive way to provide a synthetic overview of the data and serological reactivity. Multivariate analyses were performed for orthoebolavirus seropositivity as well as separately for the 3 antigens for which most reactive samples were observed. We developed Generalized Linear Mixed Models (GLMM) to test the impact of several drivers on the probability of a bat being positive by serology for orthoebolavirus antigens (lme4 package in R software). We used results of the serological tests as the response variable with a binomial distribution, and species, sex, age and season as explanatory variables with fixed effects (See Table S2 from Additional file 1). To account for clustered samples collected during the same sampling session at the same site, we included sampling period as a random effect to control for repeated measures from the same trapping session.

In order to select the best model, we used the glmulti function (Rsoftware, glmulti package (30)) to rank all possible models combining the explanatory variables by the Akaike Information Criterion (AIC). We selected the models with the lowest AIC (See Table S3 in Additional file 1). The odds ratios were estimated for each explanatory variable. We also used the emmeans and ggeffects functions to obtain estimated marginal means for the response variable and visualize the effect of each explanatory variable. Analysis were carried out on R software 4.1.2.

### PCR screening for filoviruses

Oral and rectal swabs from a subset of bats were selected according to the analysis of serological results, extracted and tested for filovirus RNA by a semi-nested PCR following the protocols described in Djomsi et al. 2022 (24). The PCR targeted a 630 bp fragment of the L gene using degenerated primers that detect a wide diversity of filoviruses, i.e., Filo-MOD-FWD: TITTYTCHVTICAAAAICAYTGGG and FiloL.conR: ACCATCATRTTRCTIGGRAAKGCTTT in round 1 and Filo-MOD-FWD: TITTYTCHVTI-CAAAAICAYTGGG, and Filo-MOD-RVS: GCYTCISMIAIIGTTTGIACATT in round 2 (7)

## Results

### Bat sampling and serological results

We analyzed blood samples collected longitudinally from 1,427 wild frugivorous bats: 196 *E. helvum*, 186 *H. monstrosus*, 380 *M. angolensis* and 665 *R. aegyptiacus* (Table 1 and Table S4 from Additional file 1). Descriptive analyses show a total of 336/1427 (24%) samples that are reactive with at least one antigen. Proportions of positive samples per bat species, sex, age class and season are presented in Table 2. A total of 24/1427 (1.7%) bats are reactive with minimum 2 antigens of a specific orthoebolavirus and are thus seropositive for at least one of the orthoebolavirus species tested for (Table 3). Orthoebolavirus seroprevalence is highest in the *R. aegyptiacus* population (3%).

**Table 1.**
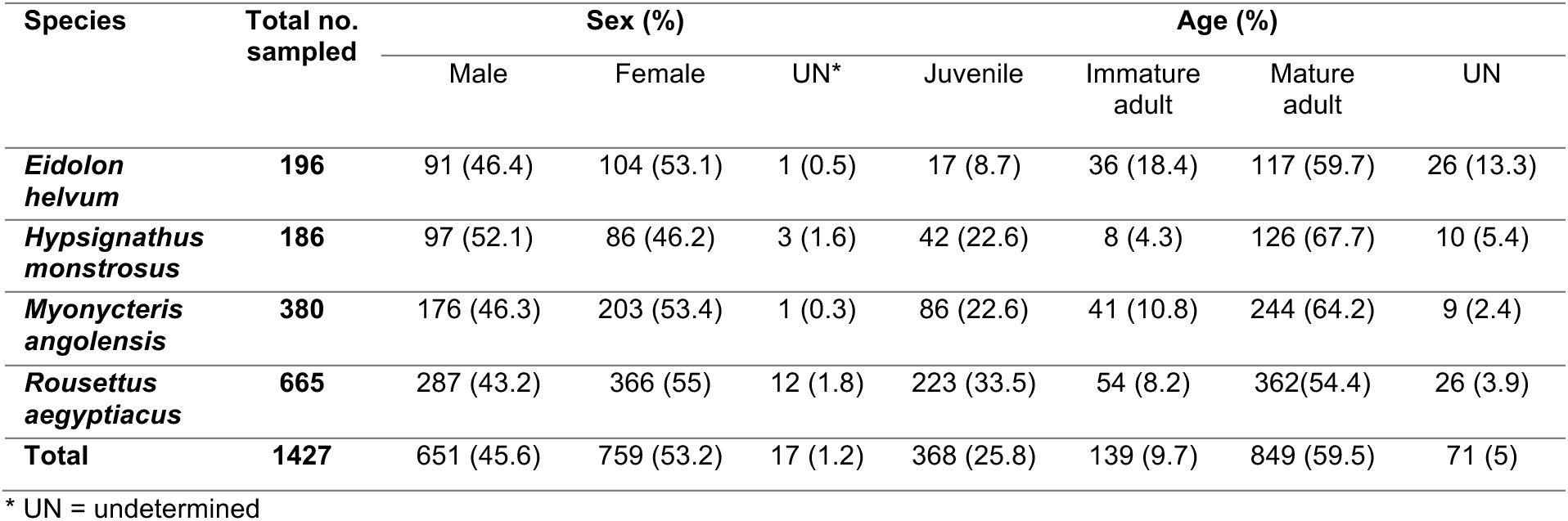
Bats sampled for ebolavirus serology by species, sex and age class.

**Table 2.** Number and percentage of samples positive for at least 1 of the ebolavirus antigens according to the M+4SD cutoff method.

| Species | Total |  |  |  |  |  |  |  |  |  |  |  |
| --- | --- | --- | --- | --- | --- | --- | --- | --- | --- | --- | --- | --- |
|  | Tot. | Pos. | % (95% CI) |  |  |  |  |  |  |  |  |  |
| *E. helvum* * | 196 | 69 | **35** (29-42) |  |  |  |  |  |  |  |  |  |
| *H. monstrosus* | 186 | 17 | **9** (6-14) |  |  |  |  |  |  |  |  |  |
| *M. angolensis* | 380 | 37 | **10** (7-13) |  |  |  |  |  |  |  |  |  |
| *R. aegyptiacus* | 665 | 213 | **32** (29-36) |  |  |  |  |  |  |  |  |  |
| Total | 1427 | 336 | **24** (21-26) |  |  |  |  |  |  |  |  |  |
| Season | Dry |  |  | Wet |  |  |  |  |  |  |  |  |
|  | Tot. | Pos. | % (95% CI) | Tot. | Pos. | % (95% CI) |  |  |  |  |  |  |
| *E. helvum* | 195 | 69 | **35** (29-42) | 1 | 0 | **0** (0-95) |  |  |  |  |  |  |
| *H. monstrosus* | 173 | 14 | **8** (5-13) | 13 | 3 | **23** (8-50) |  |  |  |  |  |  |
| *M. angolensis* | 183 | 8 | **4** (2-8) | 197 | 29 | **15** (10-20) |  |  |  |  |  |  |
| *R. aegyptiacus* | 388 | 148 | **38** (33-43) | 277 | 65 | **23** (19-29) |  |  |  |  |  |  |
| Total | 939 | 239 | **25** (23-28) | 488 | 97 | **20** (17-24) |  |  |  |  |  |  |
| Sex | Male |  |  | Female |  |  | UN |  |  |  |  |  |
|  | Tot. | Pos. | % (95% CI) | Tot. | Pos. | % (95% CI) | Tot. | Pos. | % (95% CI) |  |  |  |
| *E. helvum* | 91 | 43 | **47** (37-57) | 104 | 25 | **24** (17-33) | 1 | 1 | **100** (5-100) |  |  |  |
| *H. monstrosus* | 97 | 9 | **9** (5-17) | 86 | 8 | **9** (5-17) | 3 | 0 | **0** (0-56) |  |  |  |
| *M. angolensis* | 176 | 16 | **9** (6-14) | 203 | 21 | **10** (7-15) | 1 | 0 | **0** (0-95) |  |  |  |
| *R. aegyptiacus* | 287 | 100 | **35** (30-41) | 366 | 107 | **29** (25-34) | 12 | 6 | **50** (25-75) |  |  |  |
| Total | 651 | 168 | **26** (23-29) | 759 | 161 | **21** (18-24) | 17 | 7 | **40** (22-64) |  |  |  |
| Age | Juvenile |  |  | Immature adult |  |  | Mature adult |  |  | UN |  |  |
|  | Tot. | Pos. | % (95% CI) | Tot. | Pos. | % (95% CI) | Tot. | Pos. | % (95% CI) | Tot. | Pos. | % (95% CI) |
| *E. helvum* | 17 | 8 | **47** (26-69) | 36 | 15 | **42** (27-58) | 117 | 34 | **29** (22-38) | 26 | 12 | **46** (29-65) |
| *H. monstrosus* | 42 | 2 | **5** (1-16) | 8 | 1 | **13** (1-47) | 126 | 14 | **11** (7-18) | 10 | 0 | **0** (0-28) |
| *M. angolensis* | 86 | 7 | **8** (4-16) | 41 | 8 | **20** (10-34) | 244 | 19 | **8** (5-12) | 9 | 3 | **33** (12-65) |
| *R. aegyptiacus* | 223 | 58 | **26** (21-32) | 54 | 21 | **39** (27-52) | 362 | 121 | **33** (29-38) | 26 | 13 | **50** (32-68) |
| Total | 368 | 75 | **20** (17-25) | 139 | 45 | **32** (25-41) | 849 | 188 | **22** (19-25) | 71 | 28 | **39** (29-51) |

\* *E.* = *Eidolon*, *H.* = *Hypsignathus*, *M.* = *Myonycteris*, *R.* = *Rousettus*, Pos. = positive, Tot. = total, UN = undetermined, CI = Confidence Interval

**Table 3.** Number and percentage of bats seropositive for orthoebolaviruses (positive for minimum two antigens of the same orthoebolavirus according to the M+4SD cutoff method)

|  | EBOV |  |  | SUDV |  | BDBV |  | Seropositive for min.<br>1 orthoebolavirus |  |
| --- | --- | --- | --- | --- | --- | --- | --- | --- | --- |
|  | Tot. | Pos. | % (95% CI) | Pos. | % (95% CI) | Pos. | % (95% CI) | Pos. | % (95% CI) |
| *E. helvum* * | 196 | 0 | **0** (0-1.9) | 2 | **1** (0.3-3.6) | 0 | **0** (0-1.9) | 2 | **1** (0.3-3.6) |
| *H. monstrosus* | 186 | 0 | **0** (0-2) | 0 | **0** (0-2) | 0 | **0** (0-2) | 0 | **0** (0-2) |
| *M. angolensis* | 380 | 0 | **0** (0-1) | 2 | **0.5** (0.1-1.9) | 0 | **0** (0-1) | 2 | **0.5** (0.1-1.9) |
| *R. aegyptiacus* | 665 | 9 | **1.4** (0.7-2.6) | 13 | **2** (1.1-3.3) | 1 | **0.2** (0-0.8) | 20 | **3** (2-4.6) |
| Total | 1427 | 9 | **0.6** (0.3-1.2) | 17 | **1.2** (0.7-1.9) | 1 | **0.1** (0-0.4) | 24 | **1.7** (1.1-2.5) |

\* *E.* = *Eidolon*, *H.* = *Hypsignathus*, *M.* = *Myonycteris*, *R.* = *Rousettus*, Pos. = positive, Tot. = total, min. = minimum, CI = Confidence Interval

Regarding seroprevalences for each antigen of the different viruses, GPs were the recombinant proteins for which most reactions were observed: 9.8% of the samples showed reactivity against the EBOV Kissidougou strain, 18.8% against SUDV and 9.6% against BDBV versus 0.1 to 3.3 % against other antigens (Table 4). Lower reactivity for VP40 and NP antigens was observed for each bat species. Simultaneous reactivity to GP and NP of the same virus was rare and observed for 0.1% of the samples for EBOV and 0.6% for SUDV. The simultaneous reactivity to the same antigen of different viral lineages was frequent: 53% of the samples were reactive to GP of at least two orthoebolavirus species, 23% to VP40 and no cross-reactivity was observed for NP (See Table S5 from Additional file 1).

**Table 4.** Number (percentage) of bat blood samples reactive for each orthoebolavirus antigen according to the M+4SD cutoff method.

| Orthoebolavirus | Antigen* | *E. helvum*<br>(n=196) |  | *H. monstrosus*<br>(n=186) |  | *M. angolensis*<br>(n=380) |  | *R. aegyptiacus*<br>(n=665) |  | Total<br>(n=1427) |  |
| --- | --- | --- | --- | --- | --- | --- | --- | --- | --- | --- | --- |
|  |  | no. (%) | 95% CI | no. (%) | 95% CI | no. (%) | 95% CI | no. (%) | 95% CI | no. (%) | 95% CI |
| Ebola | NP | **1 (0.5)** | 0.03-3 | **1 (0.5)** | 0.03-3 | **1 (0.3)** | 0.01-1 | **11 (2)** | 0.9-3 | **14 (1)** | 0.6-2 |
|  | GP K | **35 (18)** | 13-24 | **5 (3)** | 1-6 | **7 (2)** | 0.9-4 | **93 (14)** | 12-17 | **140 (10)** | 8-11 |
|  | GP M | **10 (5)** | 3-9 | **0** | 0-2 | **0** | 0-1 | **37 (6)** | 4-8 | **47 (3)** | 2-4 |
|  | VP40 | **1 (0.5)** | 0.3-3 | **0** | 0-2 | **1 (0.2)** | 0.01-1 | **17 (3)** | 2-4 | **19 (1)** | 0.9-2 |
|  | NP+GP | **0** | 0-2 | **0** | 0-2 | **0** | 0-1 | **2 (0.3)** | 0.08-1 | **2 (0.1)** | 0.04-0.5 |
| Sudan | NP | **1 (0.5)** | 0.03-3 | **1 (0.5)** | 0.03-3 | **7 (2)** | 0.9-4 | **19 (3)** | 2-4 | **28 (2)** | 1-3 |
|  | GP | **66 (34)** | 27-41 | **14 (8)** | 5-12 | **18 (5)** | 3-7 | **171(26)** | 23-29 | **269 (19)** | 17-21 |
|  | VP40 | **3 (2)** | 0.5-4 | **0** | 0-2 | **3 (0.8)** | 0.3-2 | **18 (3)** | 2-4 | **24 (2)** | 1-2 |
|  | NP+GP | **1 (0.5)** | 0.03-3 | **0** | 0-2 | **1 (0.3)** | 0.01-1 | **7 (1)** | 0,5-2 | **9 (0.6)** | 0.3-1 |
| Bundibugyo | GP | **32 (16)** | 12-22 | **2 (1)** | 0.3-4 | **15 (4)** | 2-6 | **88 (13)** | 11-16 | **137 (10)** | 8-10 |
|  | VP40 | **0** | 0-2 | **0** | 0-2 | **1 (0.3)** | 0.01-2 | **7 (1)** | 0.5-2 | **8 (0.6)** | 0.3-1 |
| Reston | GP | **0** | 0-2 | **1 (0.5)** | 0.03-3 | **0** | 0-2 | **1 (0.2)** | 0.008-0.8 | **2 (0.1)** | 0.04-0.5 |

\*NP = nucleoprotein; GP = glycoproteine (K = Kissidougou strain; M = Mayinga strain); VP = Viral protein

### Factors influencing the presence of anti-orthoebolavirus antibodies in bats

We performed analysis on orthoebolavirus seropositivity, as well as on the presence of antibodies against the following 3 antigens: GP Ebola virus Kissidougou (GP EBOV-K), GP Sudan virus (GP SUDV) and GP Bundibugyo virus (GP BDBV). The GLMM was not converging for orthoebolavirus seropositivity, due to the low seroprevalence rates. However, for each separate GP antigen we obtained fairly similar best fitted models (Table 5), i.e. the Species, Sex and Age variables were consistently selected in the best models for each antigen. The variable Season was also selected in the best models for GP EBOV-K and GP BDBV and models with or without Season were near to equivalent for GP SUDV (Table 5).

**Table 5.**
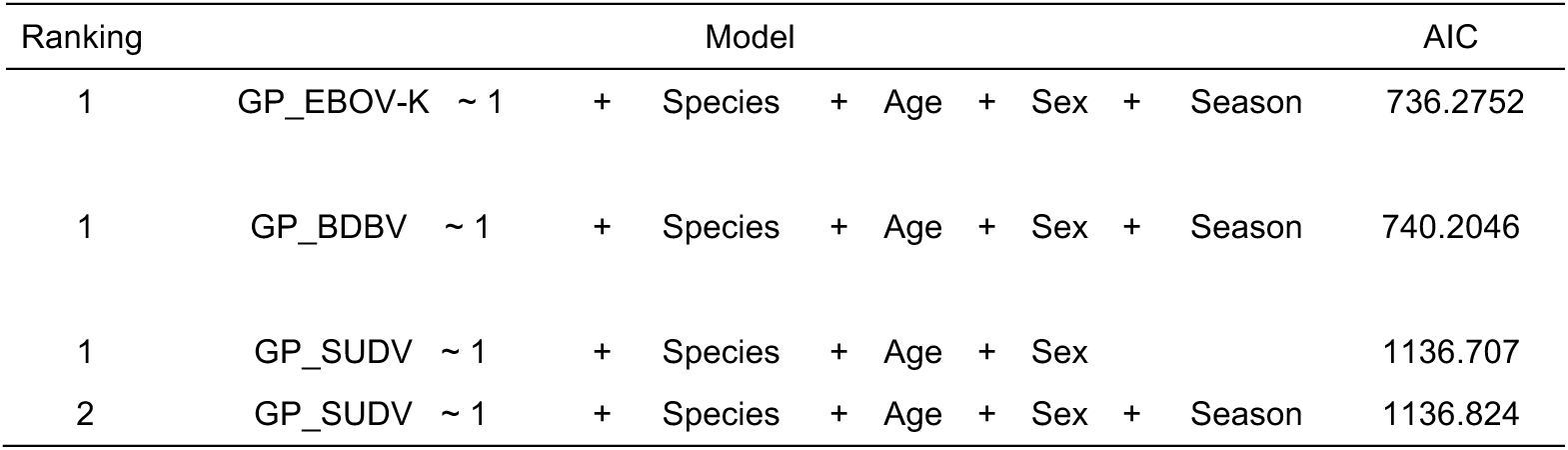
Best models obtained for the presence of antibodies against GP EBOV Kissidougou, GP BDBV and the 2 best models for GP SUDV.

Odds ratios (OR) for each explanatory variable selected in the best models are displayed with confidence intervals in Table 6. Interestingly, *H. monstrosus* and *M. angolensis* had consistently and significantly lower GP seroprevalences than *E. helvum* (OR ranging from 0.08 to 0.19 and 0.11 to 0.30 respectively, p < 0.002). Immature adults had consistently higher GP seroprevalences than juveniles in all models (OR ranging from 1.84 to 2.33), although it was only significant for GP EBOV-K and GP SUDV models (p < 0.021) but not for GP BDBV (p = 0.069). Males had consistently and significantly higher GP seroprevalences than females for all the models (OR ranging from 1.56 to 1.72, p < 0.034).

**Table 6.** Regression table representing the odds ratios of the best statistical models for GP EBOV-K, GP BDBV and GP SUDV.

| GP EBOV-K* AIC=736.3 |  |  |  | GP BDBV AIC=740.2 |  |  | GP SUDV AIC=1136.7 |  |  |
| --- | --- | --- | --- | --- | --- | --- | --- | --- | --- |
| Variables | OR | 95% CI | p-value* | OR | 95% IC | p-value* | OR | 95% IC | p-value* |
| **Species** |  |  |  |  |  |  |  |  |  |
| *E. helvum* | — | — |  | — | — |  | — | — |  |
| *H. monstrosus* | 0.18 | 0.07, 0.49 | <0.001 | 0.08 | 0.02, 0.36 | <0.001 | 0.19 | 0.10, 0.37 | <0.001 |
| *M. angolensis* | 0.13 | 0.05, 0.34 | <0.001 | 0.30 | 0.14, 0.64 | 0.002 | 0.11 | 0.06, 0.20 | <0.001 |
| *R. aegyptiacus* | 1.21 | 0.67, 2.17 | 0.5 | 1.31 | 0.73, 2.32 | 0.4 | 0.76 | 0.48, 1.21 | 0.3 |
| **Sex** |  |  |  |  |  |  |  |  |  |
| Female | — | — |  | — | — |  | — | — |  |
| Male | 1.72 | 1.15, 2.57 | 0.008 | 1.56 | 1.03, 2.36 | 0.034 | 1.70 | 1.10, 2.30 | <0.001 |
| **Age** |  |  |  |  |  |  |  |  |  |
| Juvenile | — | — |  | — | — |  | — | — |  |
| Immature | 2.33 | 1.17, 4.62 | 0.016 | 1.91 | 0.95, 3.83 | 0.069 | 1.84 | 1.10, 3.10 | 0.021 |
| Mature | 1.55 | 0.94, 2.54 | 0.085 | 1.40 | 0.82, 2.39 | 0.2 | 1.03 | 0.72, 1.48 | 0.9 |
| **Season** |  |  |  |  |  |  |  |  |  |
| Wet | — | — |  | — | — |  |  |  |  |
| Dry | 2.10 | 1.01, 4.36 | 0.046 | 1.72 | 1.02, 2.92 | 0.043 |  |  |  |

\*significant <0.05; GP = glycoprotein; EBOV-K = Ebola virus Kissidougou strain; BDBV = Bundibugyo virus; SUDV = Sudan virus;
OR = Odds Ratio; CI = Confidence Interval

Similarly, the estimated marginal means of GP seroprevalences were significantly higher for *R. aegyptiacus* and *E. helvum* compared to *H. monstrosus* and *M. angolensis* (Figure 2 and Additional file 1 Table S6) and for males compared to females (Figure 3), whatever the antigen. GP seroprevalences were significantly higher for immature adults compared to juveniles for GP EBOV-K and compared to juveniles and adults for GP SUDV (Figure 4). The GP seroprevalences were also significantly higher in the dry season compared to the rainy season for GP EBOV-K and GP BDBV (Figure 5).

**Figure 2.**
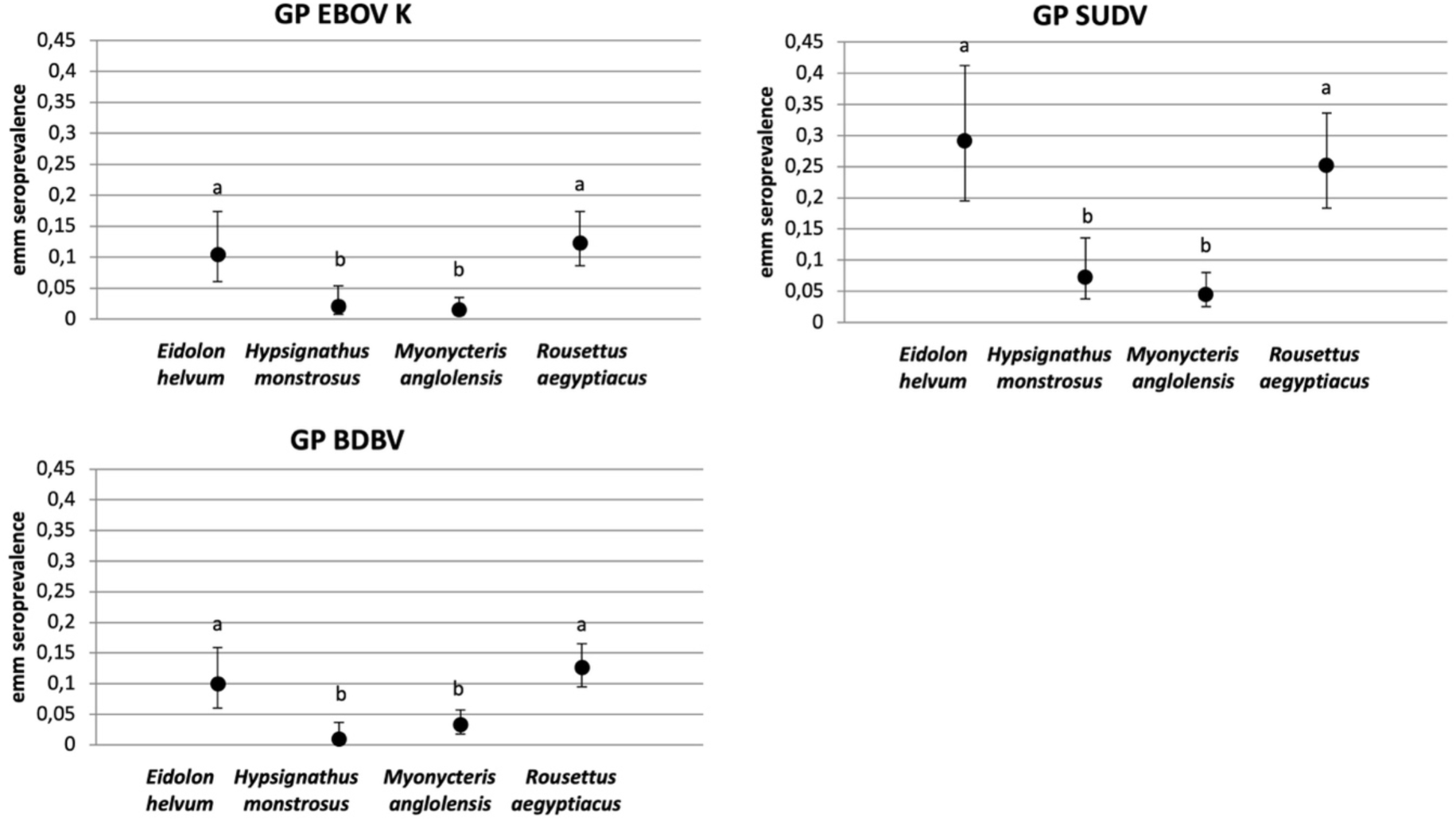
Estimated marginal means (emm) of bat seroprevalence depending on the species and the antigens (estimated marginal means annotated with different letters are significantly different (p < 0.05))

**Figure 3.**
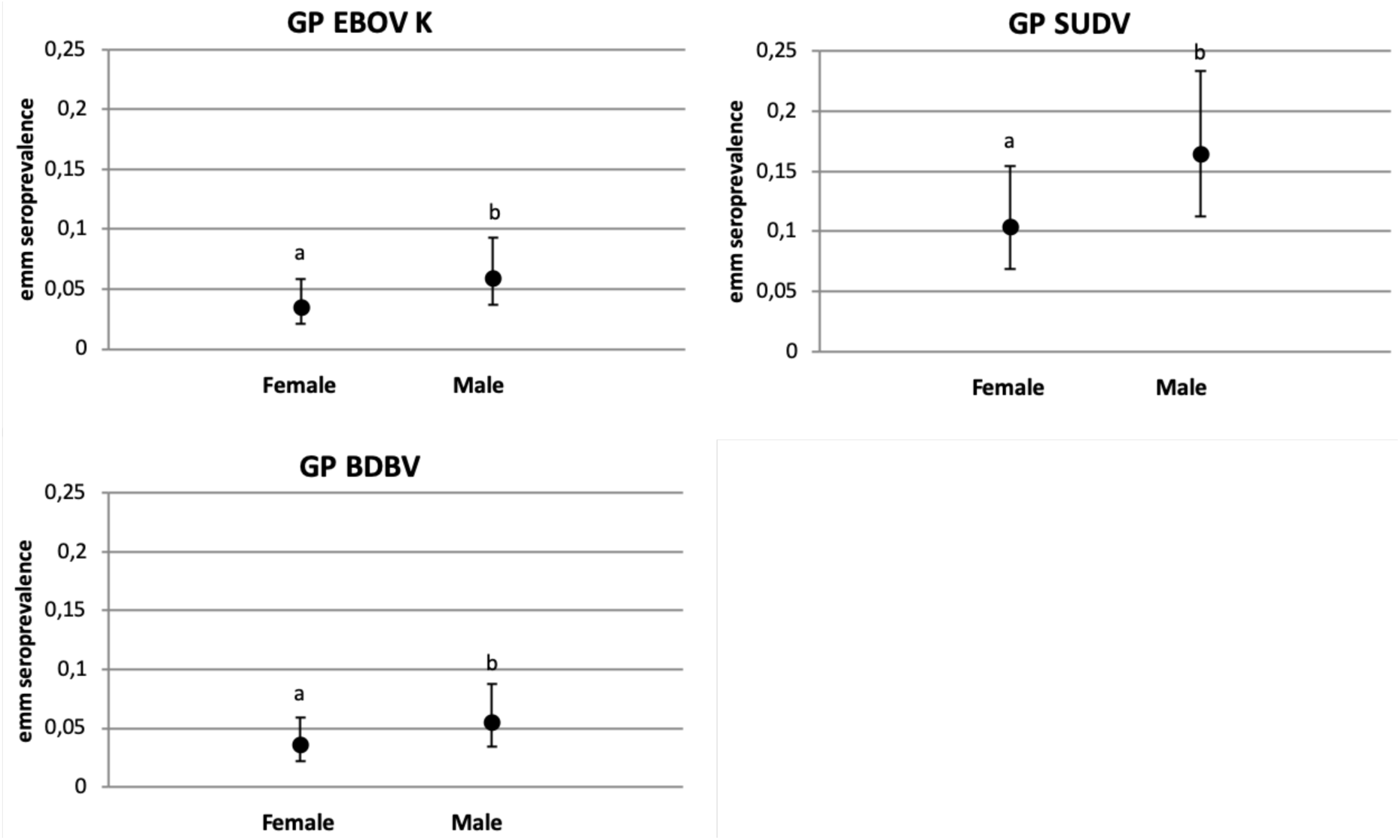
Estimated marginal means of bat seroprevalence depending on the sex and the antigens (estimated marginal means annotated with different letters are significantly different (p < 0.05))

**Figure 4.**
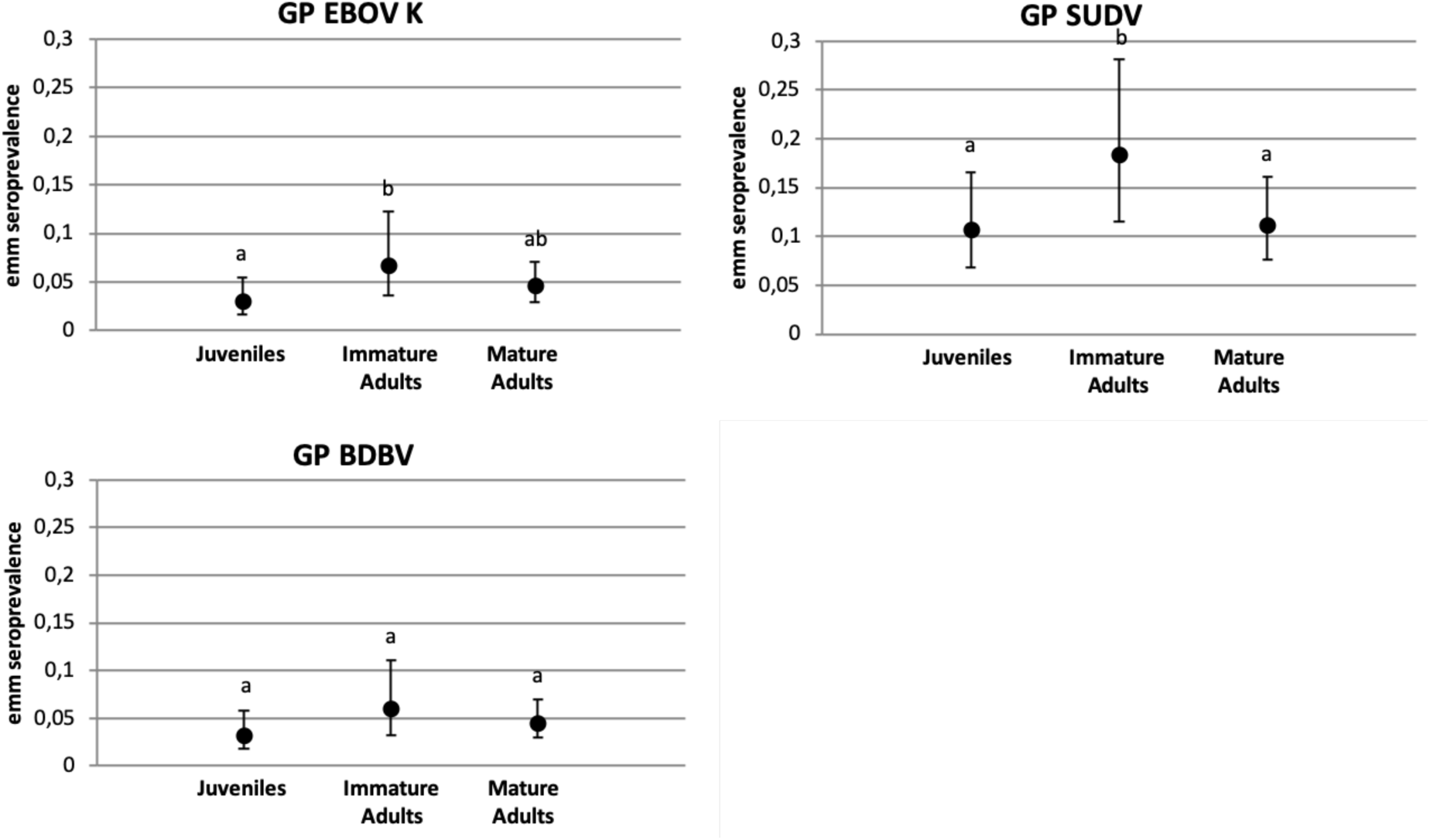
Estimated marginal means of bat seroprevalence depending on the age class and the antigens (estimated marginal means annotated with different letters are significantly different (p < 0.05))

**Figure 5.**
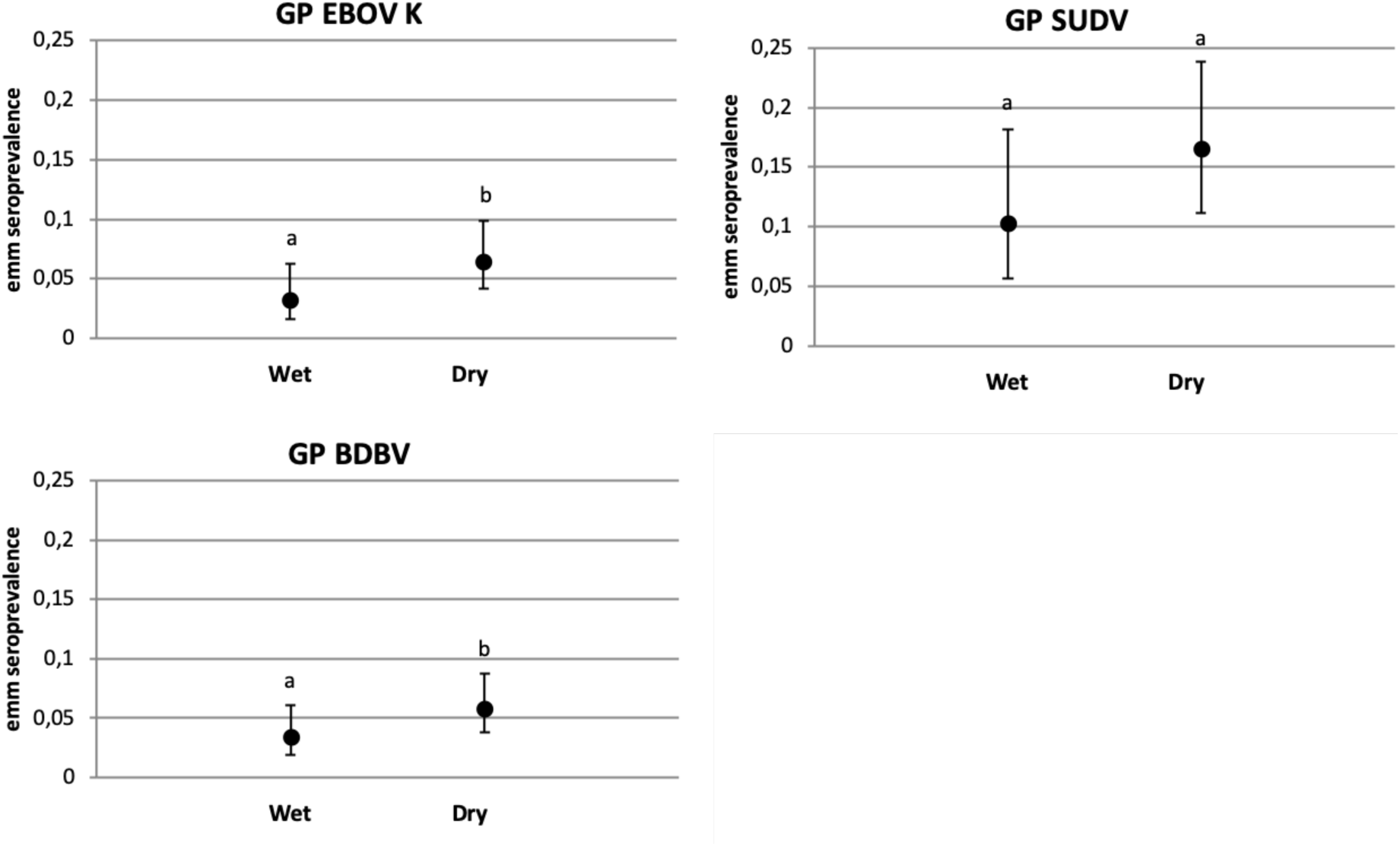
Estimated marginal means of bat seroprevalence depending on the season and the antigens (estimated marginal means annotated with different letters are significantly different (p < 0.05))

### Filovirus RNA detection

Given the results of the serological and statistical analysis, oral (195) and rectal (141) swabs from 142 *R. aegyptiacus* and 53 *E. helvum* were screened by PCR. Juveniles and immature adults, as well as a subset of adults which were serologically positive to at least one antigen were selected (see Table S7 from Additional file 1). In addition, 64 *H. monstrosus* (64 oral and 63 rectal swabs) as well as 75 *M. angolensis* (75 oral and 75 rectal swabs) were also tested given that *H. monstrosus* is one of the 3 African fruit bat species in which viral EBOV RNA has been detected in previous studies (6) and that antibodies against orthoebolaviruses were detected in both species. All samples were negative.

## Discussion

Although a number of cross-sectional studies and significant sampling efforts show the presence of antibodies against orthoebolaviruses in several bat species (8,14,28), only one reported detection of Ebola virus RNA in three fruit bats species (6). This low detection rate of viral genetic material makes it difficult to understand the dynamics of orthoebolavirus circulation in bats, the role of bats in orthoebolavirus maintenance and inter-species transmissions and related zoonotic risks. Antibody detection remains therefore a valuable indirect indicator of orthoebolavirus circulation in bats even if its interpretation remains challenging (8,31). Lack of positive controls, standardized cut-off determination methods, serological cross-reactivity, knowledge on antibody response dynamics in bats and viral molecular evidence generally complicate interpretation of results.

Here we opted for a longitudinal sampling strategy which allowed us to gain further insight into dynamics of orthoebolaviruses or closely related filoviruses in fruit bats, which is valuable for guiding further research efforts and surveillance strategies. Serological results are in line with cross-sectional studies carried out in the same region (8,32), confirming seropositivity for several orthoebolavirus antigens in the four bat species of interest, the proportion of samples reacting with at least one antigen (23,5% versus 18,2% and 18,9% for the least stringent cut-off) and simultaneous reactivity against same antigens of different orthoebolavirus species. Antibody kinetics in human EBOD survivors have been shown to vary for GP and NP antigens, with NP-antibodies declining faster (33). This might contribute to the higher seroprevalence observed here for antibodies against GP antigens, and the rarity of simultaneous reactivity to GP and NP antigens, also found in other studies (8,32). The lack of specificity due to antibody cross-reactivity between filoviruses and the use of single GP antigens per virus for statistical analysis might constitute limitations of this study. However, clear and biologically meaningful patterns are observed with the methodology used. This suggests that the serological results for the GP antigens as interpreted here do reflect past or present infection with orthoebolavirus species tested for in this study or with one or several other, perhaps not yet identified, filoviruses which induce antibodies that cross-react with known orthoebolavirus antigens.

Our results show that the presence of antibodies against orthoebolavirus GP antigens in fruit bats is best explained by bat species, sex, age as well as season. Best fitted models were almost identical regardless of the orthoebolavirus tested for (GP EBOV-K, GP SUDV or GP BDBV). We opted for interpretation of results in terms of orthoebolavirus GP in general and not specific to orthoebolavirus species as the homogeneity of the model outputs for the different orthoebolavirus GP antigens might be linked to cross-reactivity between GP antigens, as has been described in previous studies (8,29,31).

*E. helvum* and *R. aegyptiacus* consistently showed significantly higher orthoebolavirus GP seroprevalences than *H. monstrosus* and *M. angolensis* and *R. aegyptiacus* had the highest seroprevalence when taking into account minimum 2 antigens per virus. *R. aegyptiacus* are reservoir hosts of Marburg virus, a filovirus also known to circulate in West Africa (17,34), and have been experimentally shown to be less likely to maintain orthoebolaviruses (35). One explanation for our serological findings in *R. aegyptiacus* might be that they reflect Marburg virus infection through IgG cross-reactivity with orthoebolavirus antigens rather than infection with an orthoebolavirus. Serological IgG cross-reactivity between orthoebolaviruses and Marburg virus has experimentally been shown to be very limited in bats using indirect ELISA with infectious-based filovirus antigens (31). However, we used recombinant viral protein antigens for our multiplex serological assays, hence cross-reaction of anti-Marburg virus IgG might be higher compared to assays based on infectious-based antigens (31). An alternative explanation for our serological results in *R. aegytpiacus* could be regular exposure of *R. aegyptiacus* to known or unknown orthoebolaviruses through other infected animals, leading to antibody response, and potential low-level replication followed by rapid viral clearance. *R. aegyptiacus* is actually accountable for the highest prevalence of orthoebolavirus seropositive bats (minimum 2 antigens) of our study. Despite the fact that *R. aegyptiacus* might not be able to maintain known orthoebolaviruses infection, they have actually been shown to produce virus-specific IgG when challenged experimentally with orthoebolaviruses (35). Moreover, inoculation experiments on *R. aegyptiacus* use viral strains originating from human epidemics, which might have been subject to mutations during passages on cells. The extent to which other potentially circulating orthoebolaviruses are able to cause more or less persistent infection in this species is unknown. Our results for *E. helvum* are in line with other serological surveys which have detected antibodies against orthoebolavirus GP in colonies from various geographical sites (8,24,36). An experimental study has shown reduced susceptibility of Eidolon cells to EBOV (37). However, susceptibility to other filoviruses might vary. In the same way as for *R. aegyptiacus*, the detected antibody response might reflect exposure to orthoebolavirus(es) followed by limited replication and rapid viral clearance or a certain degree of infection by one or several other closely related filoviruses.

Beyond species-specific physiology and immunity, bat behaviour and ecology certainly also contribute to inter-species variability in antibody responses. Large social groups have been shown to allow higher rates of intra-species contacts which, for some pathogens, might facilitate transmission and increase prevalence (38). *E. helvum* and *R. aegyptiacus* are social species living in large colonies (tens or hundreds of thousands of individuals), therefore offering a large potential for viral transmission, while *M. angolensis* and *H. monstrosus* live in smaller groups (less than fifty for *M. angolensis* and twenty for *H. monstrosus*) (26). *H. monstrosus* is known to form leks which gather thousands of individuals during the mating season in central Africa (39), but leks seem to be smaller and sparse in forested Guinea (field observation). *E. helvum* migrate long distances and form a panmictic population across most of the African continent (40), which might also favor repeated viral reintroductions in colonies. Specific feeding and roosting habits might lead to different degrees of inter-species contacts with other bat or non-bat hosts of orthoebolaviruses and thus viral exposure. As a matter of fact, *R. aegyptiacus* and *H. monstrosus* were observed sharing the same feeding sites in Macenta (T. Pouliquen, pers. comm.), forested Guinea, hence exposure of one of these species to filoviruses hosted by the other followed by immune reaction cannot be excluded. Actually, specific Marburg antibodies have been documented in *H. monstrosus* in Gabon (41). Understanding the role of bats in orthoebolavirus maintenance and transmission requires a perspective at the community level rather than species level. Further studies on bat demographics, ecology, behaviour as well as identification of the viruses responsible for the detected antibody responses are still needed to elucidate these species differences in seroprevalence and their role in viral maintenance and transmission, particularly in Guinea.

This study suggests for the first time that male fruit bats are more likely to be seropositive against orthoebolavirus GP than females despite species-specific sampling limitations. This gender difference was however not observed in a longitudinal study on *E. helvum* in Cameroon (24). Further longitudinal species-specific studies focusing on this question are needed to investigate this in different species, and to explore the explanatory hypothesis based on observations made on humans. Studies on human populations have shown that EBOV frequently persists in male survivors’ semen, sometimes for several years as suggested by the recent epidemic recrudescence in Guinea (2,42). This might result in repeated antigenic stimulation and slower decrease of antibodies (33,43). We cannot exclude that this scenario also occurs in some bat species, perhaps at specific times, and underlies higher GP seroprevalences in males (under the assumption that antibodies decline over time when there is no persistence). Behaviour such as mating and aggressive contacts might also increase the frequency of close interactions of males with conspecifics compared to females and favor microbial transmissions, as has been observed for other host species (44). In the same line, a recent study on astro- and paramyxovirus circulation in bats has shown higher viral detection rates in males (45).

The effect of age on the probability of detecting anti-orthoebolavirus GP antibodies, with peak seroprevalences in the age class of immature adults, is in line with Marburg virus infection dynamics where viral infection peaks in older juvenile (6 months old) *R. aegyptiacus* (23) as well as with orthoebolavirus serological profiles described in *E. helvum* in Cameroun (24,32). Passive immunity, made up of maternal antibodies to Marburg, wanes in juvenile *R. aegyptiacus* aged 4 to 5 months, making them vulnerable to infection with the virus (46). This could also be the case for other filoviruses in fruit bats and explain increases in orthoebolavirus GP seroprevalence in immature adults following peaks of viral infections once older juveniles (4 to 6 months) are losing their passive immunity. Juvenile bats sampled in the present study were at least several months old as they were flying independently, which makes this hypothesis plausible in explaining the observed pattern of increased GP seroprevalence in immature adults compared to these older juveniles. Regardless of the underlying mechanisms, and similarly to what has been observed in *E. helvum* (24), results suggest that a seroconversion against orthoebolavirus or closely related filovirus takes place during the transition between juvenile and immature adults. The duration of immunity against orthoebolaviruses in bats being a knowledge gap, the lower antibody prevalence observed here in mature adults compared to immature adults, especially for GP SUDV, could suggest a waning in time of the antibody response, with potential re-stimulation in certain individuals, as suggested in humans (43).

Finally, orthoebolavirus GP seroprevalences seem to be generally higher during the dry season (November to April) compared to the rainy season. *E. helvum,* which have relatively high seroprevalence rates, are almost absent of the study site during the rainy season due to their migratory behaviour. Reported seroprevalences may thus also reflect bat population dynamics. Moreover, this seasonal pattern is most probably also related to other factors which we couldn’t measure in this study hence caution is needed with interpretation of this result. Food abundance such as fruits can be limited during the dry season. Consequently, this could lead to an exacerbated intra and interspecific competition for food (47) and increased viral transmission followed by increased seroprevalences. Moreover, bat species studied here present one or several birth pulses during this period, and for instance new *E. helvum* recruits reach about 6-7 months of age at the beginning of the dry season. Rainfall and bat reproduction are probably strongly interlinked (22,48), hence seasonal effects detected here are likely to reflect the effect of reproductive life-cycle (and thus also age) on seroprevalence. The data per bat species was insufficient to identify temporal patterns at a finer scale or assess the effect of parameters such as period of copulation, parturition, lactation and weaning for each bat species. However, other studies show that these periods can correspond to increased risks of viral transmissions or impact viral maintenance (41,49,50).

Despite important sampling efforts over time, sample size remains small in relation to the size of bat populations, and species-specific sample size was too limited to go into further analysis at the bat species level and at a finer temporal scale. Nevertheless, this study provides valuable information to guide further research on filovirus circulation in wildlife as well as surveillance in the study region. Due to potential serological cross-reactivity between orthoebolavirus species (29,31,33), serological observations may reflect past infections with various orthoebolaviruses or closely related filoviruses, including undiscovered viruses. The regular discovery of novel filovirus species across continents such as new filoviruses in China (51,52), Bombali virus in four African countries (7,53–55), Lloviu virus in three European countries (18,56,57), suggests that the full filovirus diversity has only been partly explored so far. Molecular data which could help determine the filovirus species responsible for the antibodies detected in our study and confirm serological interpretation remains limited. Future identification of the virus(es) underlying these serological signals requires sequencing of viral genetic material and virus isolation, and is essential to understand filovirus maintenance, transmission dynamics and risks of inter-species spillover or emergence. Our results suggest that intensive sampling efforts to search for evidence of and characterize active viral infection should include a strong focus on older juveniles as well as male bats. A special attention should be given to *E. helvum* and *R. aegyptiacus* which present significantly higher seroprevalences compared to the other bat species, without excluding research on other species. In the same way, surveillance of spillover events related to filovirus circulation in bats should be reinforced where and when circulation and risks might be enhanced: at the interface with bat colonies when older juveniles are highly prevalent, especially *E. helvum* (during the dry season for forested Guinea) and *R. aegyptiacus* (already demonstrated for Marburg virus in *R. aegyptiacus*). Based on these hypotheses, we screened a selection of individuals for filovirus RNA, but all samples were negative. Given the observed GP seroprevalence rates, we assume that the failure to detect filovirus RNA might be explained by either very low viral shedding at time of sampling, the circulation of unknown filoviruses which might not be detected by the PCR protocol used, or inappropriate sample type. Viral load might indeed be generally higher in internal organs than in rectal and oral swabs and it might thus be necessary to use the above-mentioned criteria to target a subset of individuals for more invasive sampling. Screening of *H. monstrosus* and *M. angolensis* which showed lower GP seroprevalence rates also revealed negative results. Nevertheless, EBOV RNA has been detected in organ samples from *H. monstrosus* in the past (6). Isolating filovirus nucleic acid from bats remains a challenge and when detected, prevalence rates based on rectal and oral swabs are often low (e.g. 1, 6 % for BOBV in Goldstein et al. 2018 (7) and 0,7% for Marburg virus in Paweska et al. 2020 (58)). Better targeted and larger sample sizes, combined with metagenomic approaches might be necessary to increase probability of filovirus RNA detection. Further experimental studies can also provide a clear insight into host competence for maintenance and transmission of known viruses and be complementary to field studies (59).

## Conclusions

This study improved the understanding of filovirus dynamics in frugivorous bats by using a longitudinal approach that helps to test the role of potentially influential factors. It emphasizes the importance of host reproductive seasonality as a driver of pathogen circulation and thus spillover risks, which should be considered when designing sampling protocols for surveillance and research. Data can for instance be used to update orthoebolavirus spillover risk maps already developed for forested Guinea (60). The study was implemented in an ecosystem exposed to intensive human transformation influencing viral ecology. Substantial sampling efforts need to be pursued and combined with data from ecological and behavioural studies on bats and inter-species relations to allow for a broader comprehension of the ecology of filoviruses at the community level. These efforts should also be replicated in different social-ecological ecosystems with different exposure to human footprint for a comparative analysis.

## List of abbreviations

AIC: Akaike Information Criterion
BDBV: Bundibugyo Ebola virus
bp: base pairs
EBOD: Ebola disease
EBOV: Ebolavirus
ELISA: Enzyme-Linked Immuno-Sorbent Assay
emm: Estimated Marginal Means
GLMM: Generalized Linear Mixed Models
GP: Glycoprotein
GP-K or GP EBOV-K: Glycoprotein Ebola virus Kissidougou strain
GP-M or GP EBOV-M: Glycoprotein Ebola virus Mayinga strain
IgG: Immunoglobulin G
MFI: Median Fluorescence Intensity
M + 4SD: mean plus four times the Standard Deviation
NP: Nucleoprotein
OR: Odds Ratios
p: p value (significant < 0.05)
PCR: Polymerase Chain Reaction
RESTV: Reston Ebola virus
RNA: Ribonucleic acid
SUDV: Sudan Ebola virus
VP40: Viral protein 40

## Declarations

### Ethics approval and consent to participate

This study was approved by the Comité National d’Ethique pour la Recherche en Santé (CNERS) in Guinea (permit number 025/CNERS/20), and the Office Guinéen des Parcs Nationaux et Réserves de Faune (OGPRF) of the Ministère de l’Environnement, et du Développement de la République de Guinée (permit 017OGUIPAR/MEEF/2017).

### Availability of data and materials

The datasets generated and analysed during the current study are available in the following repository: https://doi.org/10.18167/DVN1/KMKUQ8.

## Funding

This study was supported by the projects Ebo-Sursy (https://rr-africa.oie.int/en/projects/ebo-sursy-en/) (FOOD/2016/379–660) and BCOMING (Horizon Europe project 101059483), both funded by the European Union, and the EbOHealth project, publicly funded by I-SITE MUSE (Montpellier Université d’Excellence https://muse.edu.umontpellier.fr/en/muse-isite - MUSE2018-EbOHEALTH) through the French National Research Agency (ANR) under the “Investissements d’avenir” program (grant number ANR-16-IDEX-0006).

The funders had no role in study design, data collection and analysis, decision to publish, or preparation of the manuscript.

### Authors’ contributions

MC has contributed to the design, acquisition, analysis, interpretation of the data, and drafting of the manuscript. AC, MB have contributed to the conception, design, analysis, interpretation and revision of the manuscript. AA, MP, AKK have contributed to the design, acquisition, analysis, interpretation of the data and revision of the manuscript. AS and MID have contributed to the data acquisition, interpretation and revision of the manuscript, TP and GT have contributed to the acquisition of the data and revision of the manuscript. JC and HDN have contributed to the conception, design, data acquisition, analysis, interpretation and drafting of the manuscript. All authors read and approved the final manuscript.

## Acknowledgements

We thank the Comité National d’Ethique pour la Recherche en Santé (CNERS) and the Office Guinéen des Parcs Nationaux et Réserves de Faune (OGPRF) of the Ministère de l’Environnement, et du Développement de la République de Guinée for authorising this study. We thank the Direction Nationale des Services Vétérinaires of the Ministère de l’Agriculture et de l’Elevage of Guinea, the OGPFR, and the CERFIG for their constant support and facilitation of the project, as well as all the field staff (Aboubacar Diallo, Pierre Kabinet Kamano, Mamadou Aliou Bah, Jean Paul Guilavogui, Mamadou Aliou Sow, Aboubacar Camara Bountouraby, Aboubacar Camara, Fassou Kourouma, KoïKoï Sakouvogui).

## Additional file 1

PDF format .pdf

**Methods S1**. Molecular confirmation of bat species

**Table S1.** Mean fluorescence intensity cutoff values for different viral antigens, by orthoebolavirus lineage and statistical method used to determine cutoff according to De Nys et al, 2018.

**Table S2**. Characteristics of the variables used in the GLMM statistical models.

**Table S3**. Ranking of the top 5 best statistical models for the 3 glycoproteins by increasing AIC value carried out on Rstudio software (version 1.4.1106)

**Table S4**. Longitudinal representation of the number of bats sampled per month for orthoebolavirus serology (October 2018 - July 2020)

**Table S5**. Simultaneous reactivity of bat antibodies against antigens of different orthoebolavirus lineages (M+4SD cutoff)

**Table S6**. p-values calculated with the estimated marginal means methods to compare differences in antibody positivity according to age, sex, season and species (with the M+4SD method) (significant p-value <0.05)

**Table S7**. Samples tested by PCR for filovirus RNA

