## Additional file 1 for "Effects of biological and environmental factors on orthoebolavirus serology in bats in Guinea"

### Supporting Information

**Methods S1**. Molecular confirmation of bat species

We confirmed bat species identification recorded in the field on 19 bats by using molecular tests on Dried Blood Spots as described in Lacroix et al., 2020. We amplified an ≈800-bp fragment of mitochondrial cytochrome b using primers cytb-L140217 (forward 5' - ATGACCAACATCCGAAAATCNCAC- 3') and cytb-H15506 (reverse 5' - AGTGGRTTRGCTGGTGTRTARTTGTC -3'). To improve PCR performance for one species, we substituted the forward primer with the cytb- L2 primer which was slightly modified (C added at the 3' end) (5' - ATYTCYTCMTGATGAAAYTTYGGMTC - 3') (*1*). We purified PCR products through agarose gel (1%) and directly sequenced on an ABI 3500 sequencer (Applied Biosystems, Courtaboeuf, France). We performed BLAST (https://blast.ncbi.nlm.nih.gov/Blast.cgi) analyses to identify the most similar bat species. The samples had a query cover of 99 or 100% and a percentage identity between 98.99 and 100%. We confirmed 19 identifications recorded in the field (3 *H. monstrosus*, 8 *M. angolensis* et 8 *R. aegyptiacus*) and extrapolated to other specimens based on morphology, capture site and capture date.

**Table S1.** Mean fluorescence intensity cutoff values for different viral antigens, by Ebola virus lineage and statistical method used to determine cutoff according to De Nys et al, 2018.

| | | **Statistical method** | | |
|---|---|---|---|---|
| OBV† species, Antigen | **Mean+4SD*** | Changepoint method | Binomial method (0.01) | Exponential method (0.01) |
| **Zaire** | | | | |
| Nucleoprotein | **71** | 382 | 165 | 76 |
| Glycoprotein K | **128** | 406 | 1231 | 492 |
| Glycoprotien M | **307** | 484 | 1475 | 566 |
| Viral protein 40 | **75** | 324 | 155 | 76 |
| **Sudan** | | | | |
| Nucleoprotein | **131** | 608 | 265 | 111 |
| Glycoprotein | **251** | 929 | 2831 | 1089 |
| Viral protein 40 | **88** | 294 | 240 | 120 |
| **Bundybugyo** | | | | |
| Glycoprotein | **99** | 389 | 714 | 296 |
| Viral protein 40 | **363** | 398 | 162 | 369 |
| **Reston** | | | | |
| Glycoprotei**n** | **249** | 147 | 163 | 70 |

*Mean plus for times standard deviation.
†OBV = orthoebolavirus ; K = Kissidougou strain from West Africa 2014; M = Mayinga strain from Democratic Republic of the Congo 1976.

**Table S2**. Characteristics of the variables used in the GLMM statistical models.

| | Variables | Description | Modalities | Fixed or random effect | Hypotheses |
|---|---|---|---|---|---|
| **Response variables** | GP_EBOV-K[†]<br>GP_BDBV<br>GP_SUDV | Presence of antibodies directed against | 0 = negative<br>1 = positive * | | |
| **Explanatory variables included in different models** | **Species** | Determined according to morphological criteria** | *Eidolon helvum*<br>*Hypsygnathus monstrosus*<br>*Myonycteris angolensis*<br>*Rousettus aegyptiacus* | Fixed | Higher seroprevalence for *E. helvum* and *R. aegyptiacus* |
| | **Sex** | Determined by the sexual organs | Female<br>Male | Fixed | No difference in seroprevalence expected |
| | **Age category** | Determined by the body size, articular epiphyses and the development of sexual organs | Juvenile<br>Adult immature<br>Adult mature | Fixed | Lower seroprevalence for juveniles and higher for immature adults |
| | **Sampling period** | Corresponds to the 13 different periods of field missions | 30/10/2018 - 15/11/2018<br>02/12/2018 - 22/12/2018<br>24/01/2019 - 12/02/2019<br>04/03/2019 - 23/03/2019<br>08/04/2019 - 24/04/2019<br>21/05/2019 - 12/06/2019<br>10/07/2019 - 27/07/2019<br>17/09/2019 - 15/10/2019<br>12/11/2019 - 02/12/2019<br>15/01/2020 - 07/02/2020<br>05/05/2020 - 28/05/2020<br>19/06/2020 - 11/07/2020<br>27/11/2020 - 19/12/2020 | Random<br><br>Need to consider the non-independence of samples taken during the same sampling session | |
| | **Season** | Corresponds to the sampling season | Dry season (November-April)<br>Wet season (May-October) | Fixed | Seroprevalence higher in dry season because less resources and thus more contacts |

*higher than the cut-off M+4SD **combined to genetic confirmation for some specimen.
[†]GP = glycoprotein; EBOV-K = Kissidougou Ebola virus; SUDV = Sudan virus; BDBV = Bundibugyo virus

**Table S3**. Ranking of the top 5 best statistical models for the 3 glycoproteins by increasing AIC value carried out on Rstudio software (version 1.4.1106)

| Ranking | Response variable* | | | | Explanatory variables | | | | | | | AIC† | Weightable |
|---|---|---|---|---|---|---|---|---|---|---|---|---|---|
| 1 | GP_EBOV-K | ~ | 1 | + | Species | + | Age | + | Sex | + | Season | 736.2752 | 7.883422e-01 |
| 2 | GP_EBOV-K | ~ | 1 | + | Species | + | Age | + | Sex | | | 738.9130 | 2.108175e-01 |
| 3 | GP_EBOV-K | ~ | 1 | + | Species | + | Age | + | | | Season | 750.4477 | 6.594622e-04 |
| 4 | GP_EBOV-K | ~ | 1 | + | Species | + | Age | | | | | 753.0354 | 1.808308e-04 |
| 5 | GP_EBOV-K | ~ | 1 | + | | | Age | + | Sex | + | Season | 787.3229 | 6.483812e-12 |
| 1 | GP_BDBV | ~ | 1 | + | Species | + | Age | + | Sex | + | Season | 740.2046 | 7.933707e-01 |
| 2 | GP_BDBV | ~ | 1 | + | Species | + | Age | + | Sex | | | 742.9148 | 2.046207e-01 |
| 3 | GP_BDBV | ~ | 1 | + | Species | + | Age | + | | | Season | 752.5894 | 1.622307e-03 |
| 4 | GP_BDBV | ~ | 1 | + | Species | + | Age | | | | | 755.4596 | 3.862658e-04 |
| 5 | GP_BDBV | ~ | 1 | + | | | Age | + | Sex | + | Season | 782.4264 | 5.384180e-10 |
| 1 | GP_SUDV | ~ | 1 | + | Species | + | Age | + | Sex | | | 1136.707 | 5.145111e-01 |
| 2 | GP_SUDV | ~ | 1 | + | Species | + | Age | + | Sex | + | Season | 1136.824 | 4.854808e-01 |
| 3 | GP_SUDV | ~ | 1 | + | Species | + | Age | + | | | Season | 1160.166 | 4.143815e-06 |
| 4 | GP_SUDV | ~ | 1 | + | Species | + | Age | | | | | 1160.229 | 4.015685e-06 |
| 5 | GP_SUDV | ~ | 1 | + | Species | + | | | Sex | + | Season | 1220.866 | 2.732693e-19 |

*GP = glycorprotein; EBOV-K = Ebola virus Kissidougou strain; BDBV = Bundibugyo virus; SUDV = Sudan virus
†AIC = Akaike Information Criterion

**Table S4**. Longitudinal representation of the number of bats sampled per month for orthoebolavirus serology (October 2018 - July 2020)

| | **No. bats** | 2018 | | 2019 | | | | | | | | | | | | 2020 | | | | | | |
|---|---|---|---|---|---|---|---|---|---|---|---|---|---|---|---|---|---|---|---|---|---|---|
| | | nov | dec | jan | feb | mar | apr | may | jun | jul | aug | sep | oct | nov | dec | jan | feb | mar | apr | may | jun | jul |
| *Rousettus aegyptiacus* | **665** | 18 | 4 | 23 | 19 | 23 | 5 | 32 | 50 | 26 | 0 | 117 | 1 | 198 | 0 | 61 | 1 | 36 | 0 | 0 | 48 | 3 |
| *Myonycteris angolensis* | **380** | 5 | 22 | 2 | 6 | 14 | 4 | 34 | 20 | 67 | 0 | 21 | 17 | 21 | 0 | 67 | 0 | 42 | 0 | 0 | 19 | 19 |
| *Eidolon helvum* | **196** | 0 | 6 | 10 | 0 | 28 | 32 | 0 | 0 | 0 | 0 | 1 | 0 | 4 | 0 | 3 | 60 | 52 | 0 | 0 | 0 | 0 |
| *Hypsignathus monstrosus* | **186** | 1 | 6 | 13 | 3 | 24 | 14 | 1 | 1 | 0 | 0 | 4 | 0 | 4 | 0 | 50 | 0 | 58 | 0 | 0 | 7 | 0 |

**Table S5**. Simultaneous reactivity of bat antibodies against antigens of different orthoebolavirus species (M+4SD cutoff)

| **orthoebolaviruses antigens** | **Nbr.* (%)** |
|---|---|
| Nucleoproteins | 0/42 |
| Glycoproteins | 151/285 (52.98) |
| Viral proteins 40 | 9/39 (23.08) |

* Nbr = number of samples reacting simultaneously to the antigen of at least two different ebolavirus lineages / total number of samples reacting to the antigen.

**Table S6**. p-values calculated with the estimated marginal means methods to compare differences in antibody positivity according to age, sex, season and species (with the M+4SD method) (significant p-value <0.05)

| | GP EBOV K* | GP SUDV | GP BDBV |
|---|---|---|---|
| ***Age*** | | | |
| Juveniles-Immatures | **0.0422** | **0.0483** | 0.1633 |
| Immatures-Adults | 0.3535 | **0.0358** | 0.5769 |
| Adults-Juveniles | 0.1973 | 0.9675 | 0.3661 |
| ***Sex*** | | | |
| Male-Female | **0.0083** | **0.0008** | **0.0339** |
| ***Season*** | | | |
| Dry-Wet | **0.0456** | 0.1620 | **0.0429** |
| ***Species*** | | | |
| *E. helvum - H. monstrosus* | **0.0043** | **<.0001** | **0.0051** |
| *E. helvum - M. angolensis* | **0.0001** | **<.0001** | **0.0106** |
| *E. helvum - R. aegyptiacus* | 0.9202 | 0.8395 | 0.8009 |
| *H. monstrosus - M. angolensis* | 0.9568 | 0.5646 | 0.3478 |
| *H. monstrosus - R. aegyptiacus* | **0.0006** | **<.0001** | **0.0009** |
| *LM angolensis - R.aegyptiacus* | **<.0001** | **<.0001** | **<.0001** |

*GP = glycorprotein; EBOV-K = Ebola virus Kissidougou strain; BDBV = Bundibugyo virus; SUDV = Sudan virus

**Table S7**. Samples tested by PCR for filovirus RNA

| Species (number of bats) | Sample type | Number of samples | *Age* | | | *Sex* | | |
|---|---|---|---|---|---|---|---|---|
| | | | Juvenile | Immature adult | Mature adult | Male | Female | UN |
| *Eidolon helvum (53)* | Oral swabs | **53** | 15 | 35 | 3 | 25 | 27 | 1 |
| | Rectal swabs | **53** | 15 | 35 | 3 | 25 | 27 | 1 |
| *Rousettus aegyptiacus (142)* | Oral swabs | **142** | 89 | 24 | 29 | 72 | 67 | 3 |
| | Rectal swabs | **88** | 63 | 19 | 6 | 46 | 39 | 3 |
| Total | | **336** | | | | | | |
